# Flying honeybees adjust their reaction times to enable group cohesion

**DOI:** 10.1101/2022.06.03.494769

**Authors:** Md. Saiful Islam, Imraan A. Faruque

## Abstract

Flying insects routinely demonstrate coordinated flight in groups. How they achieve this with very limited communication, vision, and neural systems remains an open question. We measured the visual reaction time in flying honeybees while they chased a moving target, and compared in-flight reaction times for solo animals with those flying in groups. Across 425 insects, the solo honeybees show diverse reaction times (an average of 30ms and a standard deviation of 50ms). The reaction times in groups are significantly more uniform (an average of 15ms and a standard deviation of only 7ms), indicating that honeybees in group flight adjust their reaction times to match their neighbors. To investigate the role of this adjustment, we curve fit the reaction time distributions and analyzed them in a mathematical model of swarming, finding that the reaction time increases the stable region of a cohesive swarm. To verify the stabilizing effect was not an artifact of curve fitting, we then inserted the measured delays in a swarm simulation, which breaks apart under the solo reaction times and achieves stable formations for the group reaction times. Together, our findings highlight how flying animals can synchronize their reaction times in group flights to improve group cohesion.

## Reaction time in swarms

A significant barrier to creating adaptive swarms of small scale unmanned aerial systems and other challenging robotic swarms remains the provision of fast, computationally-lightweight sensing and feedback structures to support relative navigation in dynamic groups [1–3]. Insects flying in groups can serve as model systems for resource-constrained feedback on these swarming micro air vehicles [4]. Group behavior implies that complicated interactions may be used to make decisions among themselves [5–7]. Midge swarms regulate themselves relative to a dynamic moving stimulus which reveals possible interactions present in their common activity [8]. The delay associated with a swarm’s overall motion may have a dependency on number of agents in mosquitoes tracking a moving marker [9]. Neuronal networks with delays can have a significant role on the resulting biological behaviors, including generating oscillatory motions and periodic signals [10–12]. While experimental research is beginning to understand the need to quantify internal delays during insect sensing and feedback, these previous studies are limited to reporting a single average delay across all animals and do not yet account for the diversity of delays across the population or the effect of such reaction time delays on neighbor-coordinated behaviors.

## Bees as model systems and their behavior

Applying stimuli to swarms can result in swarm and individual-level responses [13–15]. The effects of environmental stimuli examining insect flight behavior and motions during visually-dominated behaviors like obstacle avoidance, landing on a wall or proboscis, and flower tracking have previously focused most on the role of ambient and external illumination levels [16–18]. Bees show collective behaviors and individual responses quantified by a hierarchical approach [19]; reasons and consequences for individual level behavioral variation in foraging honeybees were explored [20]. Excellent collective behaviors serving a group interest, such as migration, foraging, and predatory sensing and defence can be seen in honeybees, Apis mellifera L., throughout their life span [16, 21, 22].

## Bee reaction times in tracking tasks

We previously quantified the visual reaction time seen in flying insects tracking a moving light in a solitary task [23]. This laboratory experiment found that honeybees’ reaction times were diverse, varying from 4 to 115ms. The identified reaction times were measured at the individual insect level, and probability distributions were experimentally quantified for the measured delays. We used theoretical swarm communications analysis and simulations of a swarm responding to neighboring animals including these delays, which indicated that the swarm level effect of the varying reaction time is to damage the cohesion of swarm motions. This experiment was conducted in a laboratory environment using captured bees, which might influence them not to perform their normal behavior. Would an outdoor more naturalistic experiment give the same reaction time variation?

## Laboratory vs outdoor behaviors

To test the hypothesis that the reaction times may be different outdoors, we designed a new experiment to record visual tracking trajectories outdoors. The tool divides into two parts: visual stimulus and tracking system. The stimulus is a horizontally moving beehive entrance (Fig. 1 and Section. 1.1). The trajectories are recorded by a multi-camera tracking system described in method (Section. 1.1).

**Fig. 1:**
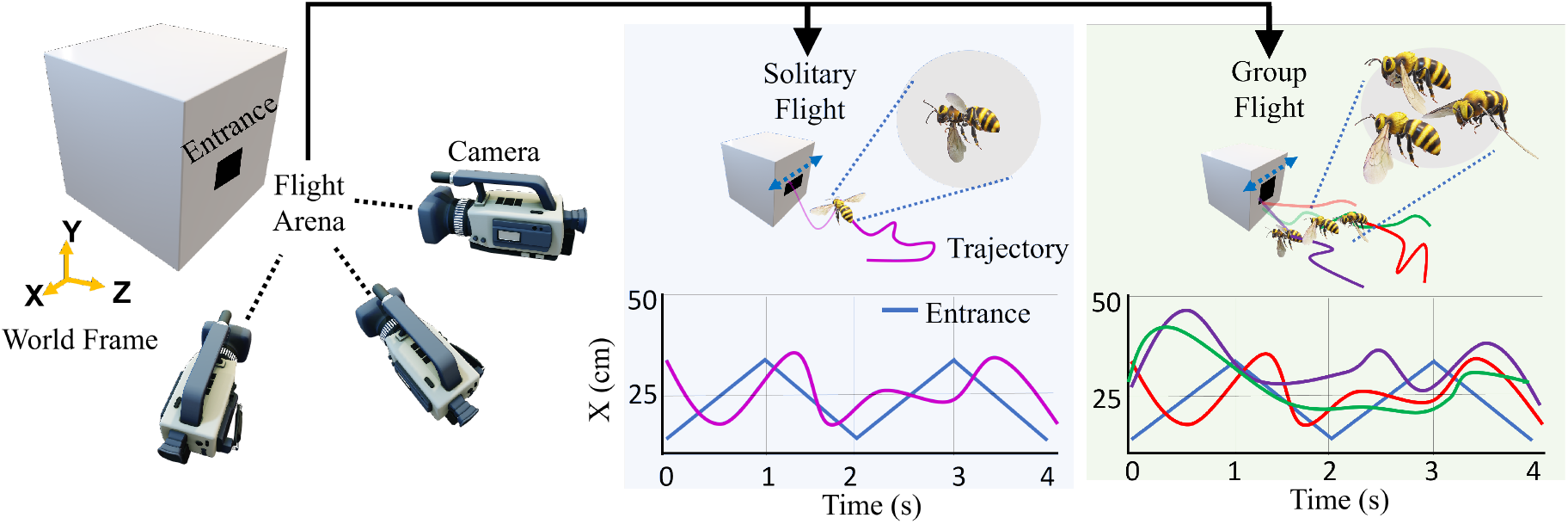
Experimental design. Experimental design consists of three camera based tracking system which record the trajectories of insects approaching to the entrance. Two types of bees’ data were segmented: solitary flight (light blue shading) and group flight (light green shading). X coordinates of entrance (blue) and insect from their 3D trajectories are considered as input and output for system identification approach, respectively.

We segmented insect’s trajectories while they approached and entered the moving entrance. We identify systems in both the time and frequency domains between the movement stimulus and the insect position. Frequency domain identification is accomplished when record lengths are sufficient (>1.5sec) and time domain when records are shorter. 105 solo insect frequency domain identifications were conducted. A representative example is shown in camera and 3D views in Figs. 2(b–d) and in the supplementary S1 Video.

**Fig. 2:**
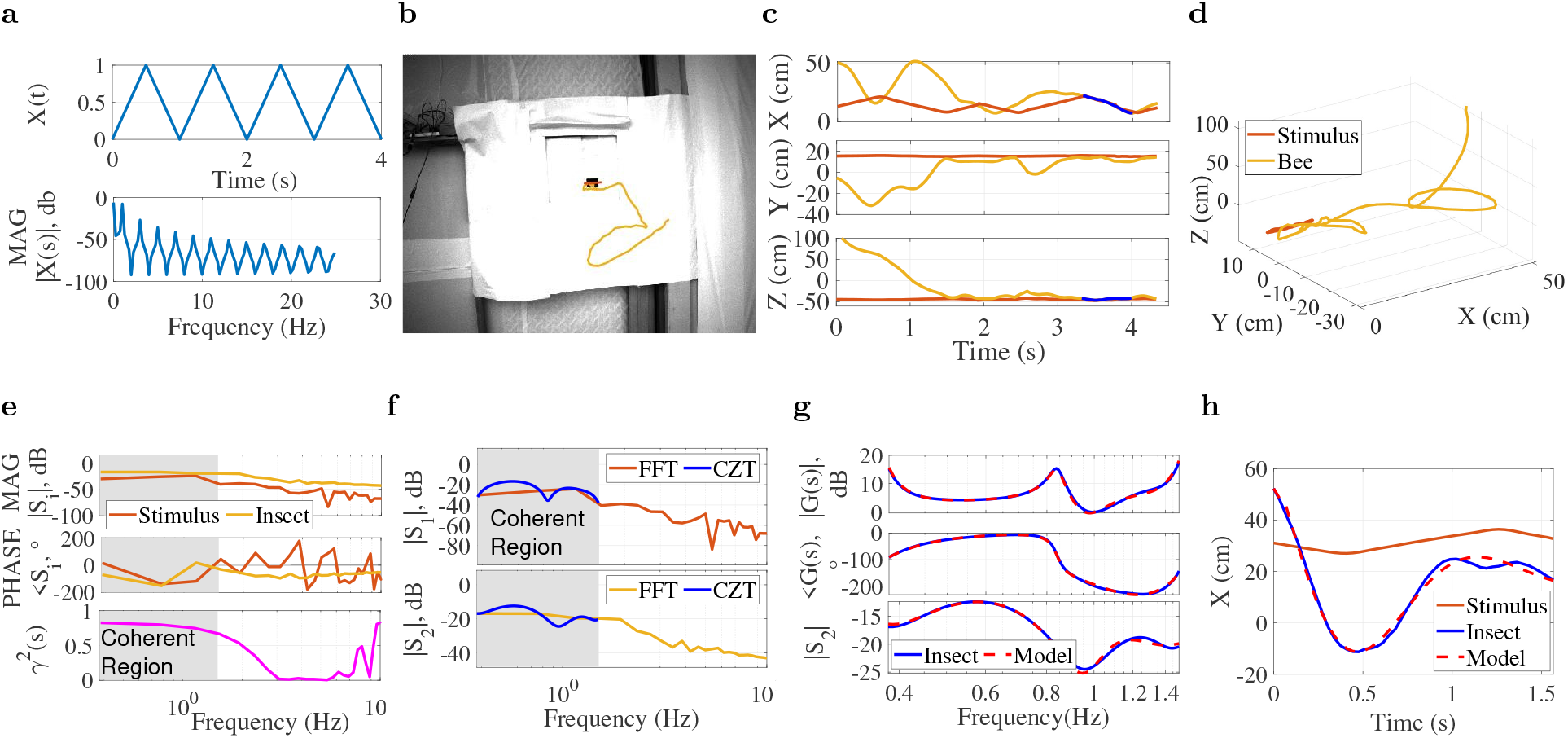
Frequency and time domain system identification examples. **a**, A triangular signal was used to apply horizontal entrance movement along the X axis of the world frame. **b**, A solitary insect’s trajectory (length 4 sec) entering the moving entrance is captured by camera-1. **c & d**, Reconstructed position coordinates and 3D trajectory for both input stimulus and insect (blue color indicates Kalman filter). **e**, Magnitude, phase, and coherence *γ*^2^ plots of the input stimulus *S*_1_ and output (insect) position *S*_2_. A linear relationship between stimulus and insect position is indicated in the shaded region below 1.5 Hz where *γ*^2^ > 0.6. **f**, Comparison of Fourier and Chirp Z-transform magnitudes for stimulus *S*_1_ and insect *S*_2_ in the region of highest coherence. **g**, The identified transfer function *G*_*e*_(*s*) (dashed red) shows strong agreement with the measured frequency response *G*(*s*) (blue) in both magnitude and phase, as do true and identified model output |*S*_2_|. **h**, Example of time domain identifications used for trials ≤ 1.5 sec show tracking performance.

Frequency transformed stimulus and output signals *S*_1_ and *S*_2_ (respectively) are used to construct a frequency response function, shown in magnitude and phase components for both fast Fourier transform (FFT) and chirp Z transforms (CZT) in Fig. 2e (see Section. 1.2.1). A coherent region of response below 1.15Hz is visible, indicating that the input and output have a strong linear relationship (as quantified by *γ*^2^ > 0.6) in this range. Some deviation from ideal tracking (0dB magnitude, 0^°^ phase) is visible in Fig. 2e, with gain showing some overshoot and a negative phase (lag) indicating a reaction time.The CZT transform was used to improve resolution in the strongly coherent range (Fig. 2f) below 1.15Hz and the CZT-derived frequency response function *G*_*CZT*_ (*s*) used to identify the equivalent transfer function. The fit error (see fit criteria at method section) statistics in Table. 1 indicate a 3 pole, 2 zero transfer function with 15 ms reaction time (transport delay) is the best fit for this example trajectory. The identified model is 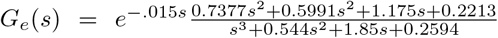, and a comparison of this identified model and experimental response functions in Fig. 2g show the strong agreement indicated by Table 1. The measured and frequency domain modeled outputs *S*_2_(*ω*) show good agreement in Fig. 2g. For short trajectories (≤ 1.5 sec), time domain system identification was performed. Fig. 2h shows an example time domain system identification that identified a model 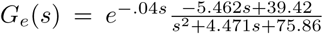 with a fit percentage of 88.13% and FPE of 1.12× 10^−3^.

**Table 1:**
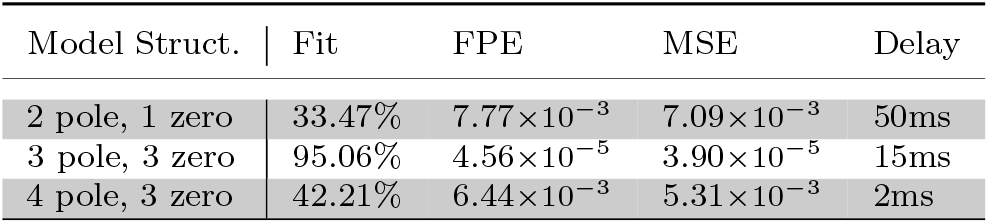
Model candidates and performance for an example insect.

## Reaction times in outdoor solitary tasks

We analysed 175 outdoor solitary trajectories as summarized in Table 2. The identified reaction times in Fig. 4(b) show that again, solitary insects have diverse reaction times, varying from 7ms to 120ms, thus the move to from laboratory to the outdoor foraging return environment retains the reaction time variation.

**Table 2:** Overall Fit Statistics.

| Role | Frequency domain |  |  | Time domain |  |  |
| --- | --- | --- | --- | --- | --- | --- |
|  | Trials | Avg. | Std. | Trials | Avg. | Std. |
| Solo | 105 | 84% | 8% | 70 | 81% | 16% |
| Group | 112 | 80% | 11% | 138 | 81% | 8% |

## Group behaviors and reaction times

When the bees tracked the entry in high traffic conditions, several bees often track the entrance at the same time. Since this group flight is likely to be visually-regulated, we asked, do insects flying in such outdoor groups still maintain the same reaction time distributions? An large variation in reaction times could be challenging for interactions in theoretical swarms, and we hypothesized that when bees fly in groups they may adjust their reaction time by interacting with other agents in group. To test for reaction time adjustments, we then identified individual insect dynamics during group approaches to the entrance using the same frequency and time domain identification process. An example of group behavior is shown in Figs. 3(a–b) and in the S2 Video. In this example, insect-1 and insect-2 are identified as a group due to their high coherence (*γ* > 0.7 up to 5 Hz). Analyzing 250 group behavior trajectories (summarized in Table 2) with an average length of 2.5 seconds showed that insects flying in the group setting demonstrate a different reaction time distribution (Fig. 4c) than those in solitary environments. In particular, the 18ms mean and significantly narrower 8ms standard deviation shows that the insects have adjusted their reaction times to reduce the diversity.

**Fig. 3:**
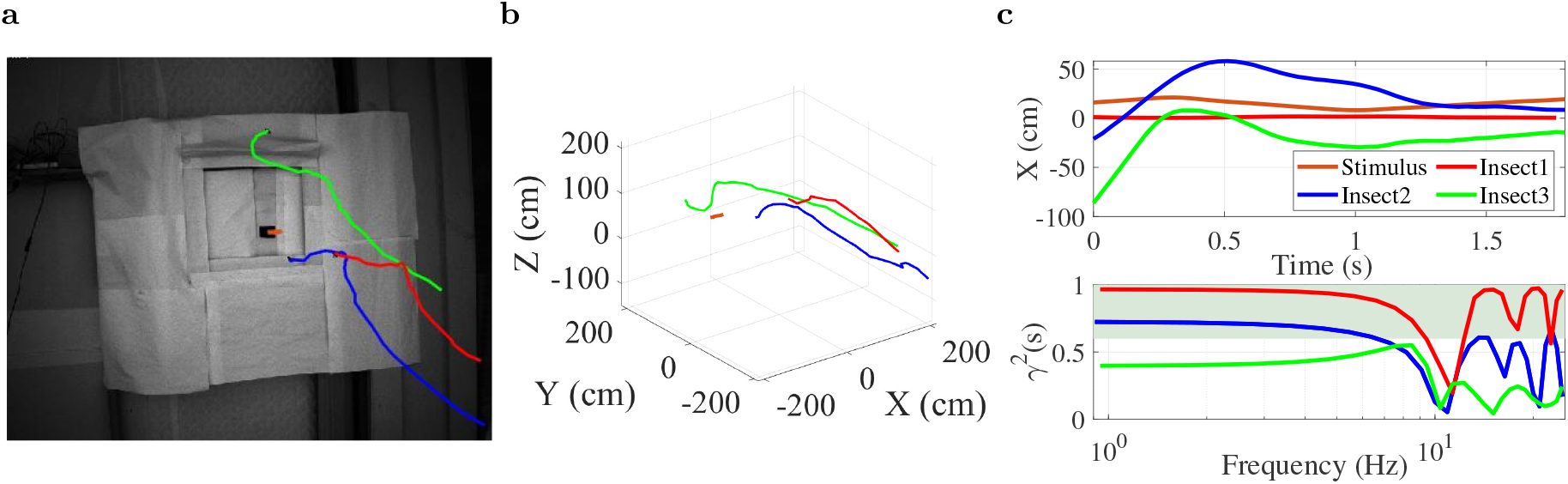
Segmentation of group insects by coherence. **a**, Three insect flight trajectories in entrance approach as seen by camera-1, **b**, Reconstructed 3D flight paths and stimulus, **c**, Insect flight trajectories and coherence. Insect1 and Insect2 are identified as group insects due to coherence *γ*^2^ > 0.7 (shaded area). Insect3’s coherence below 0.6 indicates it is not participating in group tracking.

**Fig. 4:**
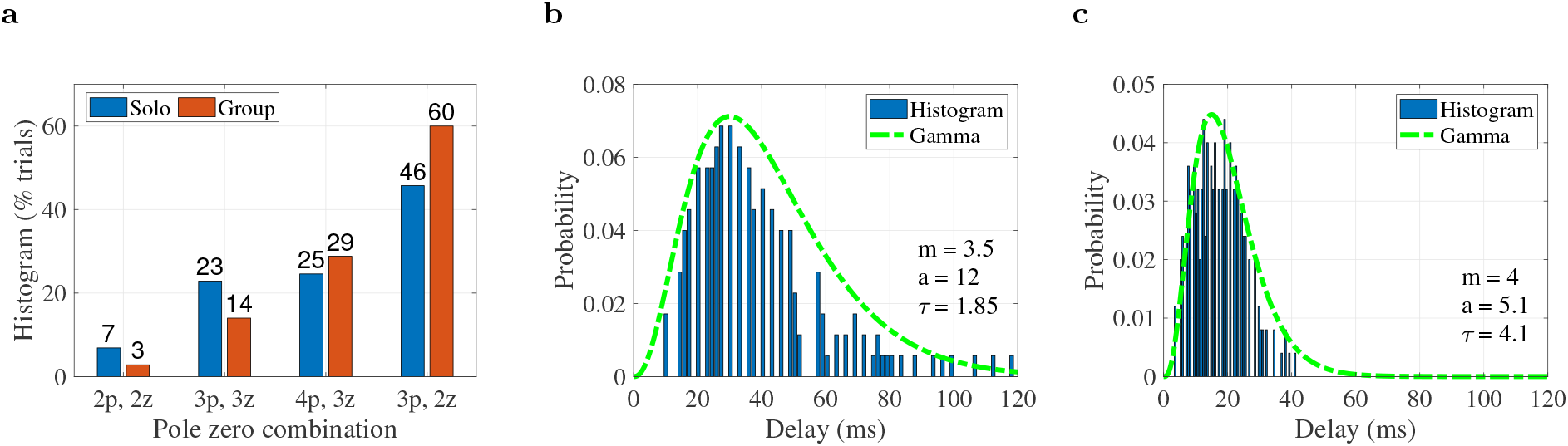
**a**, Distribution of identified model structures, as indicated by number of poles and zeros, **b**, histogram of solo insect reaction times vary from 0 to 120 ms and normalized gamma distribution, **c**, histogram and fitted distribution curve in group bees, containing a narrower band of reaction times from 0 to 45 ms.

The difference in reaction times in solo and group motions shown in Figs. 4(b–c) show that visual reaction time is adjusted from solitary to group task environments. This result motivates a theoretical question: Would this adjustment support visually-guided swarm motions, or present a challenge to such behaviors?

## Interpreting the reaction time adjustment

To provide a theoretical understanding of the impact of this experimental result, we constructed a visually interconnected swarm model with agents experiencing visual processing delays. The model is a first order dynamic system model with *N* agents experiencing delayed interconnections with each other. We built upon previous models with constant delays [24] to develop a swarm model with heterogeneous delays and weighting factors (Methods Sec. 1.3). A functional description of the reaction times is helpful for theoretical analysis, and we represented them with a gamma distribution having shape parameters *m, a*, and *τ* as indicated in Fig. 4. In this swarm model, an agent *i* updates both its position *x*_*i*_ based on its previous position and the influence of neighbors. To ensure the reaction time analysis was robust to changing preferences, we allowed the weights on these behaviors–an egocentric weight *α* and a neighbor influence *β*–to be free. We then applied mean field and bifurcation analysis on the swarm center to understand its behavior as parametric equations of *α* and *β* (Eqns. (15) and (16) in Section. 1.4). We used the experimental gamma distributions and parametric equations to find stable and unstable regions of the swarm model. The stability regions in Fig. 5a show that for low values of both egocentric and neighbor influence *α* and *β*, the swarm remains stable regardless of reaction time distribution, and high values of neighbor attraction *β* are similarly destabilizing. However, the boundary of swarm stability when using the group delays (red curve) shows a higher tolerance for neighbor attraction than the solitary delay distribution (blue curve). This finding suggests that one function of the reaction time adjustment seen in group interacting insects is to support cohesion by improving the margin of destabilization. *ω* indicates the swarm center’s oscillation frequency at the stability boundary, and shows that the delay adjustment also affects the oscillation frequency. In both cases, oscillation frequency on the stability boundary increases with egocentric influence. Group delays show higher oscillation frequencies, indicating that group reaction times support a higher frequency motion before destabilizing. In many of the recorded experiments, individual trajectory oscillations were observable; current work quantifying the trajectories indicate they are below the limit frequencies in Fig. 5a, again suggesting the insects remain below the stability boundary.

**Fig. 5:**
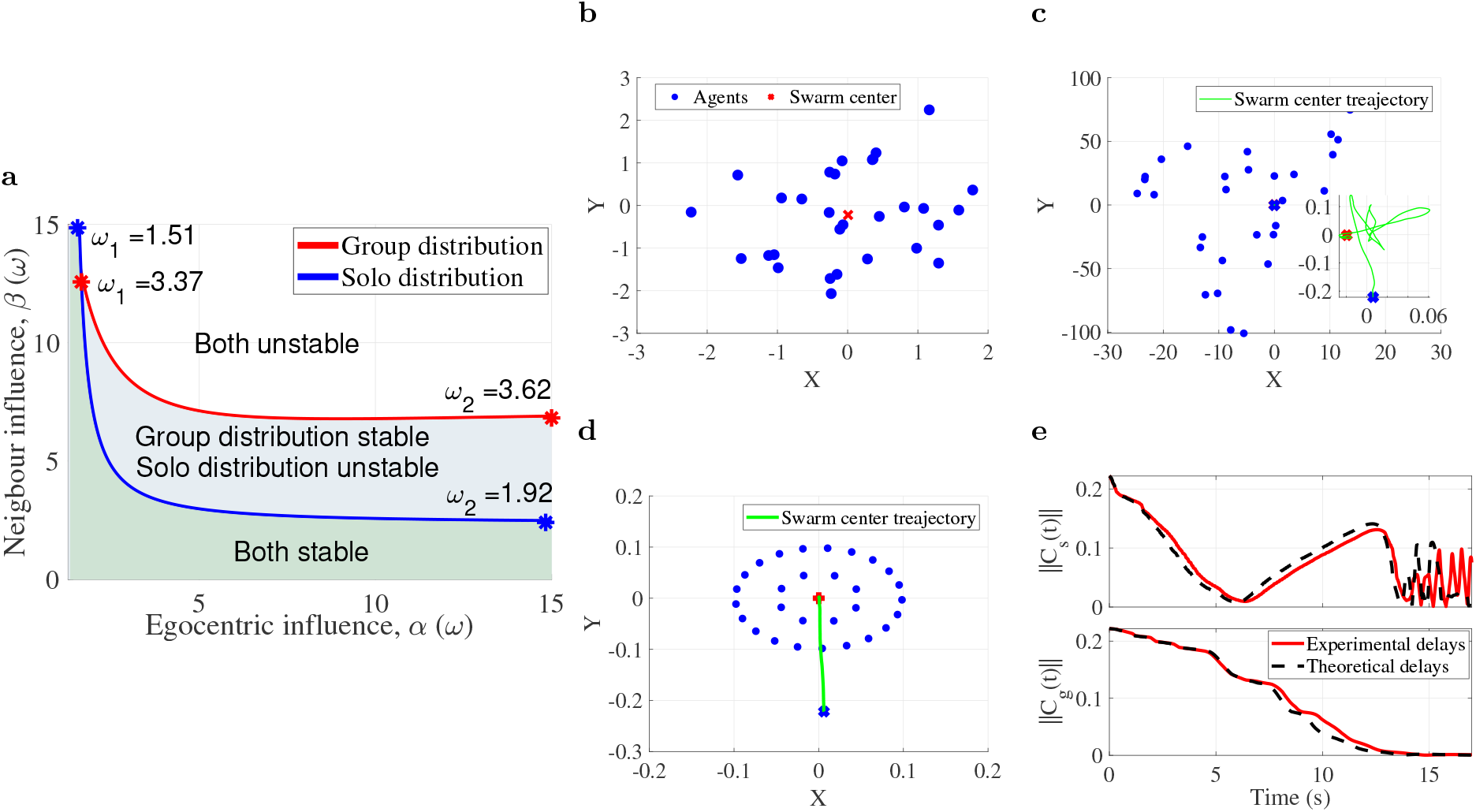
Swarm stability regions and simulations with experimental reaction times for solo and group behaviors. **a**, Stable and unstable regions for the two reaction time distributions (solo and group) were discovered by applying Eqns.(32) and (33) describing the stability boundary in terms of an individual agent’s ego influence *α* and neighbor influence *β*. Low neighbor influence factors lead to stable swarm center positions (green region) for either case. As neighbor influence grows, the swarm using a solo distribution loses stability (blue region) before the swarm using group behavior delays loses stability (white region), indicating that the group distribution provides a larger range of response weights (gains) that lead to a stable swarm. As the egocentric weight *α* grows on the stability boundary, the transition oscillation frequency *ω* rises. **b**, Random initial positions of 30 agents, **c**, ending position of agents using experimental solo reaction times, **d**, ending position of agents using experimental group reaction times, **e**, time histories of swarm center *C*_*s*_(*t*) and *C*_*g*_(*t*) norms for solo and group delays, respectively.

## Simulation verification

Some approximation is provided by the swarm model and the curve fit to a gamma distribution, and while the theoretical analysis can assess barycenter stability, it cannot yet predict formation. To explore the validity of the swarm model and examine the effects of the experimental reaction times, we simulated an interconnected visual swarm with both experimentally-quantified delays and those drawn from representative gamma distributions, beginning from the randomly distributed initial positions in Fig. 5b.

Simulations using the reaction times measured in solo and group conditions (ie, the measurements in Fig. 4) are shown in Figs. 5(b–e) for 30 agents. The time history of swarm center position in Fig. 5e shows that the swarm center stability failure seen in theoretical analysis persists in the case of experimental reaction times and is not an artifact of curve fitting. Additionally, the simulations show that the instability extends to formation, with group-measured reaction times stabilizing to a formation in Fig. 5d and solo-measured delays leading to a motion with no observable formation in Fig. 5c.

In this study, insects were recorded while tracking a moving visual stimulus in solo and group behaviors and the reaction time in their closed loop tracking was measured. We used a real-time camera-based tracking system that quantified both moving target and insect position in three dimensions, applied and system identification tools to identify the closed loop tracking dynamics between stimulus motion and insect body motion, separating the effects of open loop plant (locomotion) physics from neural reaction time. The measured insect reaction times were used to discover that reaction times vary across population. Significantly more variation (50ms standard deviation) was found when an insect was the only animal tracking the target, relative to group behaviors in which multiple insects tracked the target (8ms standard deviation). To understand the implications of the measured reaction times on visually-guided swarms, we then integrated the measured delays into a visually interacting swarm model. Analysis on this model indicates conditions needed for the center of mass’s position and allows us to map the stable and unstable regions as a function of behavior. Simulations were conducted using theoretical fits to the delays (gamma distribution) and the experimental delays. The analysis and simulation indicate that the reaction times measured in solitary conditions show unstable swarm behavior and that the group delays provide stable center of mass and cluster shape, and predict the speed of swarm center oscillations at transition.

Overall, this study quantifies reaction times in solo and group tasks and connects these measurements to theoretical limits on the allowable delays for insects in visually guided swarms. The results suggest that insects operating in group contexts adjust their delays to support swarm cohesion. This consistency between experimental measurements of solo and group tasks in flying insects with theoretical and simulated frameworks quantifying constraints is an important outcome for understanding how flying insects support swarming motions despite tight constraints on sensing and feedback. The results provide a foundation for swarming aerial robotics with limited computational resources. For these systems, in which processing delays are significant, knowledge of how animals adjust their reaction times to support stable motions will guide engineers in distributing processing tasks appropriately.

## Supplementary information

## Acknowledgments

This work was supported in part by ONR Young Investigator Award N00014-19-1-2216.

## 1 Methods

### 1.1 Experimental setup and Tracker

Figure. 6a depicts the overall study, including experimental work, theoretical analysis, and simulations. The measurement system consists of three cameras (Flir mono) with angular separations from 50 to 90 degrees. Honeybees were recorded entering and exiting a hive entrance actuated to move horizontally according to triangular frequencies. The entrance stimulus was a 2-inch square hole (tunnel entrance) that could be moved horizontally left and right using triangular frequencies along the world frame’s *X* axis. This setup included two stepper motors (Nema 23) and a motor driver (a4988), which was controlled by a micro-controller (Arduino Uno).

**Fig. 6:**
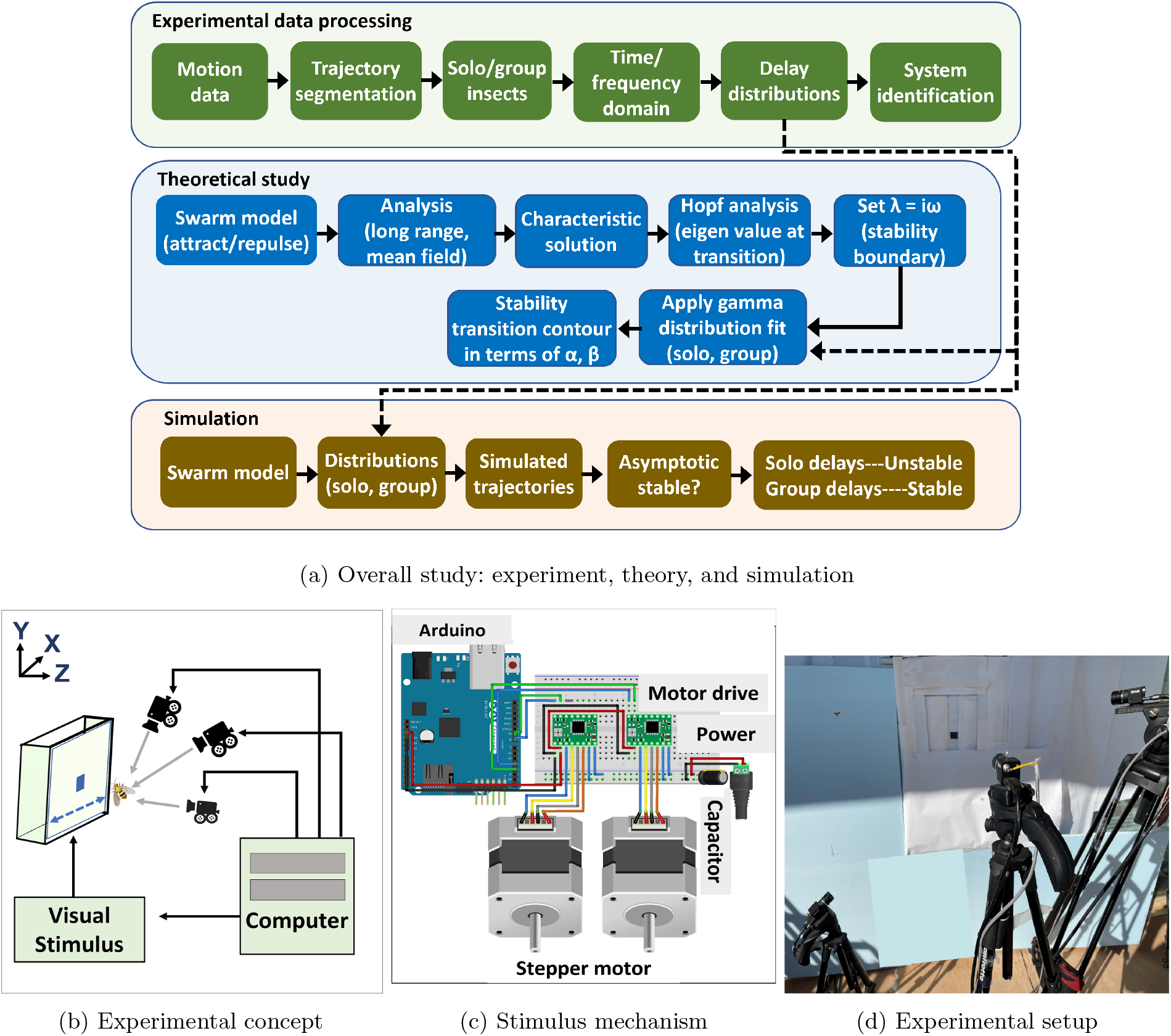
(a) Setup incorporates three components: a multi agent tracking experiment, theoretical analysis, and simulated performance. A three camera-based tracking system (b) was used to record flight paths of multiple insects (and stimulus) tracking a moving hive entrance actuated by an Arduino micro-controller and stepper motors (c), mounted in an outdoor environment (d).

The VISIONS [23] tracking system was used to record insect trajectories during solitary and group flight conditions. This study updates the tracker to track multiple insects. The functions in the tracking system are shown in Fig. 7a. To expand tracking from prior single agent tracking with VISIONS to multiple insects, we incorporated data association to match corresponding positions of each camera. The functions in the association algorithm are shown in Fig. 7b.

**Fig. 7:**
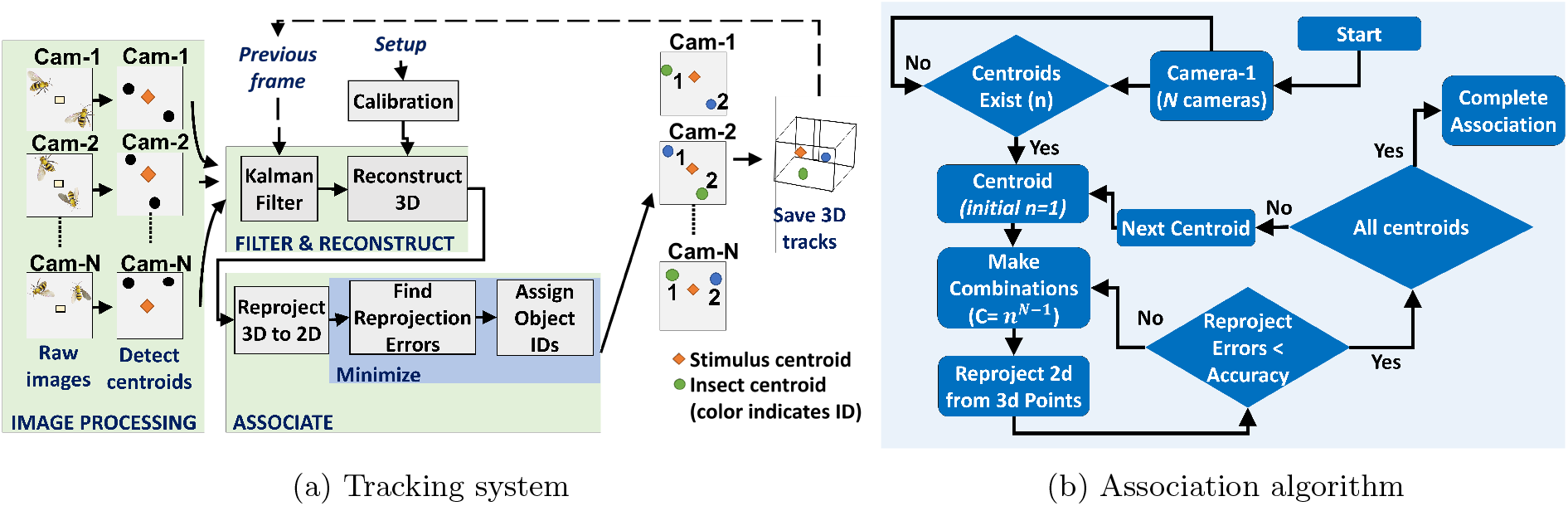
Flowchart of tracking and association algorithms for VISIONS multi-agent tracker.

The intrinsic and extrinsic parameters of the cameras were calibrated via bundle adjustment [25]. For each insect, we calculate reconstructed 3D points from all possible combinations of 2D points and then reproject them from 3D to 2D. We use the intended match from this error list if the re-projection error (difference between observed and re-projected 2d points) is less than the desired accuracy. A set of *m* insects’ 2D positions of camera-1 at time *t* is described as 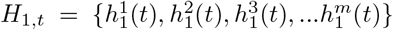. Between camera-2 and camera-*N*_*c*_, the possible combinations for each insect in *H*_1,*t*_ is 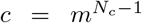. The re-projected 2D points vector is *D*_*j*_ = {*d*_1_, *d*_2_, *d*_3_, ….., *d*_*j*_}, when *j* = 1…*c*. The associated agent is taken to be

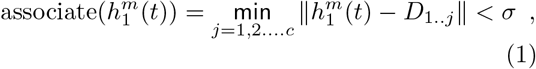

where *σ* is the desired accuracy.

### 1.2 System identification

#### 1.2.1 Frequency domain identification

When the trajectory had sufficient frequency content (trial length), we applied the identification approach for the transfer function *G*(*s*) of the insect’s position response to stimulus motion via Chirp Z transform (CZT) developed in our prior work [23] and illustrated in Fig. 8. In the frequency domain, frequency response of target tracking is described by gain, phase, and coherence. Coherence *γ*^2^(*s*) is calculated from the spectral and cross-spectral densities of input and output signals as

**Fig. 8:**
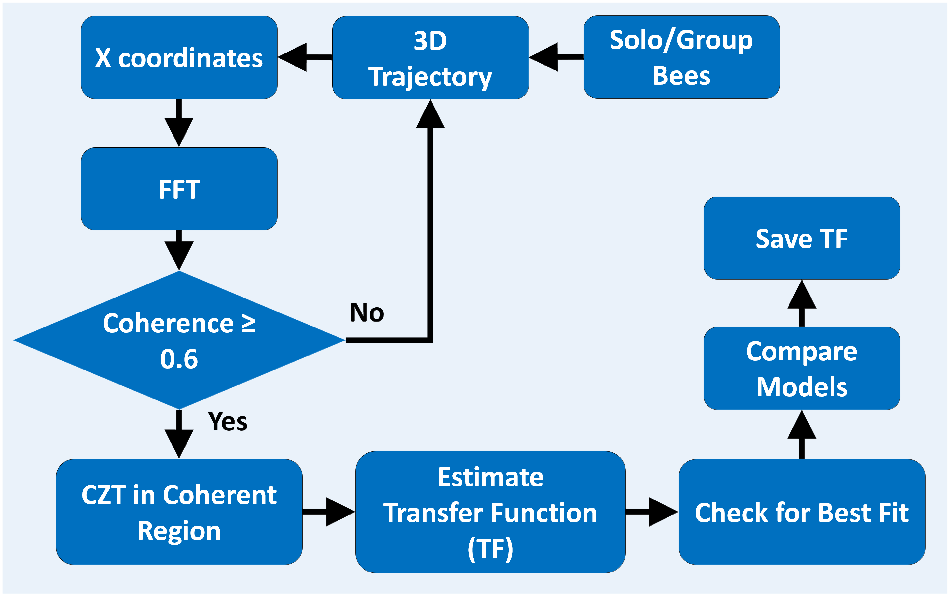
Frequency-domain system identification flowchart.

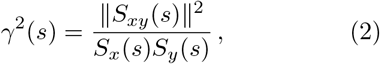

where *S*_*xy*_(*s*) denotes the cross spectral power density and *S*_*x*_(*s*) and *S*_*y*_(*s*) represent the auto power spectral density of the stimulus (input) and bee (output) coordinates, respectively. The coherence between the stimulus and insect trajectories was used to determine the linear connection throughout the frequency range. We use the frequency response function *G*(*s*) obtained from CZT transform by reference to a transfer function with an internal delay time *τ* to examine the flight dynamics and visuomotor delay of the tracking behavior. We conduct the system identification technique across several possible estimated transfer functions *G*_*e*_(*s*) and varying time delays *τ*_*i*_ ∈ [0, 200] ms. The identified transfer function minimizes the absolute difference between measured frequency response and estimated transfer functions over the region of coherence as

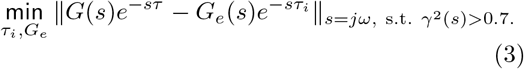

Three fit criteria FIT, MSE (mean square error) and FPE (final prediction error) are used to determine the best dynamics model [23].

#### 1.2.2 Time domain identification

For the time domain system identification, normalized least square estimation is used to find the dynamic system [26]. The discrete time domain representation of the stimulus and insect are *x*_1_(*k*) and *x*_2_(*k*), and by considering a unknown transfer function it can be written as

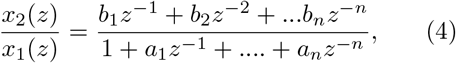

where the unknown coefficients are *a*_*i*_, *b*_*i*_; *i* = 1…*n*. The time domain solution can be written as

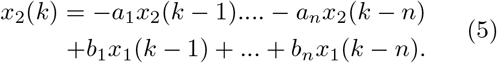

Then we construct a parametric model, *x*_2_(*k*) = *ϕ*^*T*^ (*k*)*θ*^*^ where

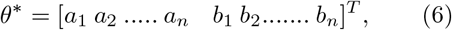

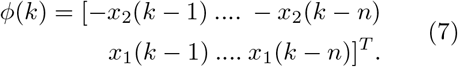

The cost function is written as

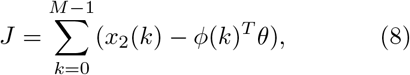

where *k* = 0 *M*. The solution 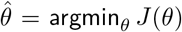 is

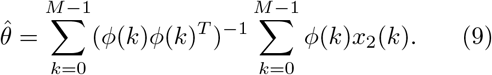

The fit criterion is defined as

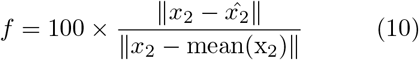

where 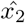 is the predicted output.

### 1.3 Mean field analysis

The swarm model is

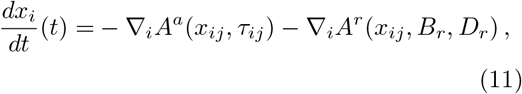

where *x*_*i*_ is (vector) position of the agent *i*,∇∇_*i*_*A*^*a*^(*x*_*ij*_, *τ*_*ij*_) and ∇_*i*_*A*^*r*^(*x*_*ij*_, *B*_*r*_, *D*_*r*_) are attractive and repulsive potentials respectively, specified as

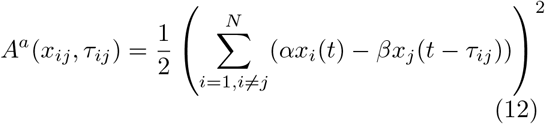

and

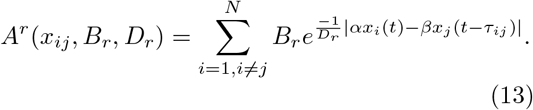

Here, *τ*_*ij*_ is the delay from agent *i* to agent *j, α* is the egocentric influence weight and *β* is the neighbor influence weight. *B*_*r*_ and *D*_*r*_ are the amplitude and applied distance of repulsive potential. At long range, stability may be determined by only the attractive potential and conversely at short range, stability may be determined by considering only the repulsive potential [23, 27]. So, for the mean field analysis we consider only the attraction potential. The center of mass of the swarm can be defined as a vector *C*(*t*) as

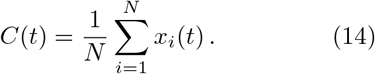

To find the position stability of the swarm, the norm of swarm center is found as 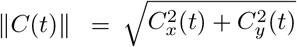, where *C*_*x*_, *C*_*y*_ are the *X* and *Y* coordinates of the swarm center respectively. Swarm center stability is then ∥*C*∥→ constant as time *t* →∇∞. Using mean field analysis (see method section) we get two coupling equations we may describe the stability contour as a function of egocentric influence *α* and neighbor influence *β* as

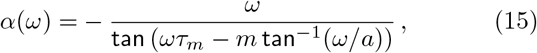

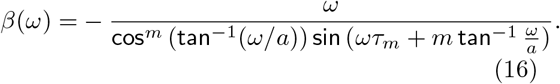

Equations (32) and (33) allow one to map the stability transition contour for a given distribution (e.g., specified *E* and *τ*_*m*_ parameters), as a function of egocentric weight *α* and neighbour influence weight *β* curves as a function of frequency *ω*.

Each agent is updated as

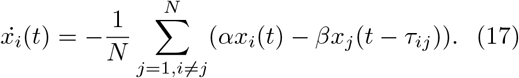

The position of each agent *x*_*i*_(*t*) = *C*(*t*) + *δx*_*i*_(*t*) and *δx*_*i*_ is the deviation from the center of mass *C*. The position derivative 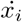 is written as

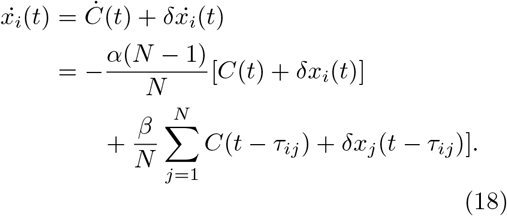

Summing over *i* and taking 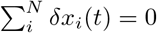,

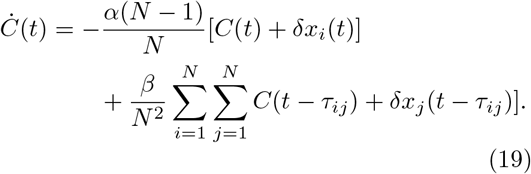

We can obtain the center of mass by approximation considering a double sum [28]. The discrete terms can be approximated by a distributed density function *g*_*τ*_ (*τ*). We can extend the discrete delays towards a distributed density function such as

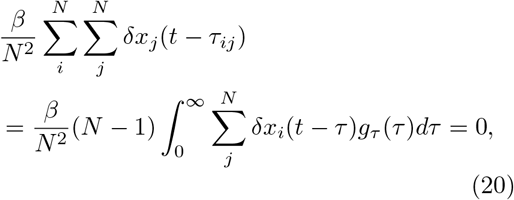

with the same approximation made for the center of mass *C*(*t*). For large number of agents 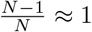. Finally, we can write the swarm center dynamics as

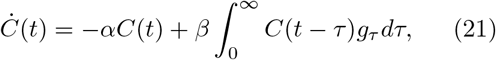

which is now a general differential equation in the form

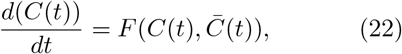

where 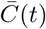 is a delay weighted state given by

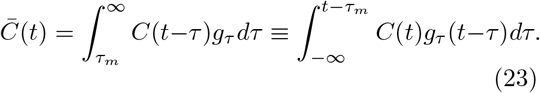

Here, *τ*_*m*_ is the minimal delay and *g*_*τ*_ is the probability density function of the distribution, such that 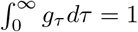.

### 1.4 Hopf bifurcation

We want to examine the sensitivity of the swarm’s local stability to changes in distribution shape. The gamma distribution has a relation between its mean with the shape. To examine local stability we need to linearize the system at a steady state solution. By taking *C*(*t*) = *ce*^*λt*^ in Eqn. (22) we obtain the characteristic solution

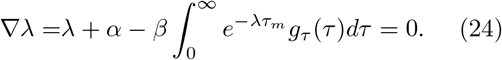

We want to examine how delay affects stability, and apply the fact that the stable/unstable transition takes place when the characteristic equation has a root with zero real part. The density of the gamma distribution is

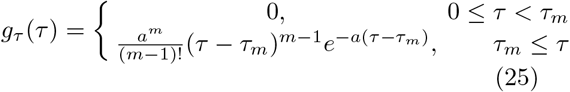

The unshifted density of the distribution can be written as 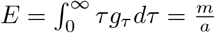, where parameters (*a, m*) specify the shape of the gamma distribution. The gamma distribution’s variance is 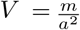, and its Laplace transform is

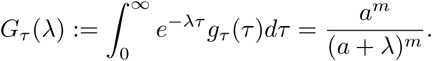

Equation (24) may then be expressed as

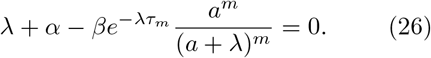

Linear systems lose stability when the roots of the characteristic equation cross the imaginary axis from left to right. To study the Hopf bifurcation, we take *λ* = *iω* and 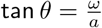 to get

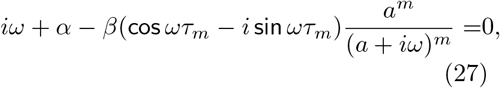

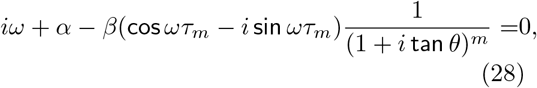

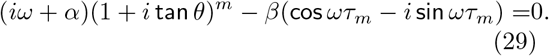

Applying de Moivre’s theorem [29] and splitting the real and imaginary parts we obtain

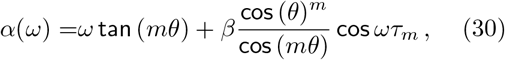

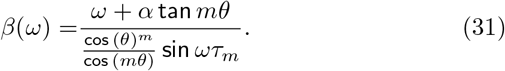

Finally, coupling the above two equations we may describe the stability contour as a function of egocentric influence *α* and neighbor influence *β* as

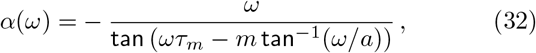

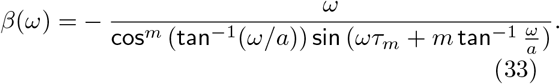

### 1.5 Simulation

Swarm simulations implemented Eqn. (11), keeping the the swarm model parameters constant as seen in Table 3. Simulations then varied the shape of the gamma distribution as in Table 3 or directly used the measured delays.

**Table 3:** Simulations used the same swarm model and different gamma distributions.

| Environment | Swarm model |  |  |  | Gamma |  |
| --- | --- | --- | --- | --- | --- | --- |
| | $\alpha$ | $\beta$ | $B_r$ | $D_r$ | $m/a$ | $\tau$ |
| Solo | 7 | 6 | 1/2 | 1 | 0.3 | 1.85 |
| Group | 7 | 6 | 1/2 | 1 | 0.78 | 4.1 |

## Appendix A Recordings and flight trajectories

***S1 Video***.

**Solo behavior**. Solitary insects track the entrance in this video.

***S2 Video***.

**Group behavior**. A subset of the 10 insects in this video track the moving entrance.

## References

[1] Schranz, M. et al. Swarm intelligence and cyber-physical systems: concepts, challenges and future trends. Swarm and Evolutionary Computation 60, 100762 (2021).

[2] Campion, M., Ranganathan, P. & Faruque, S. Uav swarm communication and control architectures: a review. Journal of Unmanned Vehicle Systems 7 (2), 93–106 (2018).

[3] Jaffe, J. S. et al. A swarm of autonomous miniature underwater robot drifters for exploring submesoscale ocean dynamics. Nature Communications 8 (1), 14189 (2017). URL https://doi.org/10.1038/ncomms14189. https://doi.org/10.1038/ncomms14189.

[4] Billah, M. A. & Faruque, I. A. Bioinspired visuomotor feedback in a multiagent group/swarm context. IEEE Transactions on Robotics 37 (2), 603–614 (2020).

[5] Berdahl, A., Torney, C. J., Ioannou, C. C., Faria, J. J. & Couzin, I. D. Emergent sensing of complex environments by mobile animal groups. Science 339 (6119), 574–576 (2013). https://doi.org/10.1126/science.1225883.

[6] Krause, J., Ruxton, G. D. & Krause, S. Swarm intelligence in animals and humans. Trends in Ecology & Evolution 25 (1), 28–34 (2010). URL https://www.sciencedirect.com/science/article/pii/S0169534709002298. https://doi.org/https://doi.org/10.1016/j.tree.2009.06.016.

[7] Downe, A. E. R. & Caspary, V. G. The swarming behaviour of chironomus riparius (diptera: Chironomidae) in the laboratory. The Canadian Entomologist 105 (1), 165–171 (1973). https://doi.org/10.4039/Ent105165-1.

[8] van der Vaart, K., Sinhuber, M., Reynolds, A. M. & Ouellette, N. T. Mechanical spectroscopy of insect swarms. Science Advances 5 (7), eaaw9305 (2019). https://doi.org/10.1126/sciadv.aaw9305.

[9] Jain, P., Singh, O. P. & Butail, S. Dynamics of mosquito swarms over a moving marker (2020). 2007.04254.

[10] Lewis, J. Autoinhibition with transcriptional delay: A simple mechanism for the zebrafish somitogenesis oscillator. Current Biology 13 (16), 1398–1408 (2003). https://doi.org/https://doi.org/10.1016/S0960-9822(03)00534-7.

[11] Morelli, L. G. et al. Delayed coupling theory of vertebrate segmentation. HFSP journal 3 (1), 55–66 (2009). URL https://doi.org/10.2976/1.3027088.

[12] MacDonald, N. Biological delay systems: Linear stability theory. Acta Applicandae Mathematica 18 (3), 297–300 (1990). https://doi.org/10.1007/BF00049132.

[13] Swain, D. T., Couzin, I. D. & Ehrich Leonard, N. Real-time feedback-controlled robotic fish for behavioral experiments with fish schools. Proceedings of the IEEE 100 (1), 150–163 (2012). https://doi.org/10.1109/JPROC.2011.2165449.

[14] Ni, R. & Ouellette, N. T. On the tensile strength of insect swarms. Physical Biology 13 (4), 045002 (2016). https://doi.org/10.1088/1478-3975/13/4/045002.

[15] Tennenbaum, M., Liu, Z., Hu, D. & Fernandez-Nieves, A. Mechanics of fire ant aggregations. Nature Materials 15 (1), 54–59 (2016). URL https://doi.org/10.1038/nmat4450. https://doi.org/10.1038/nmat4450.

[16] Baird, E., Srinivasan, M. V., Zhang, S. & Cowling, A. Visual control of flight speed in honeybees. Journal of Experimental Biology 208 (20), 3895–3905 (2005). https://doi.org/10.1242/jeb.01818.

[17] Fry, S. N., Rohrseitz, N., Straw, A. D. & Dickinson, M. H. Visual control of flight speed in Drosophila melanogaster. Journal of Experimental Biology 212 (8), 1120–1130 (2009). https://doi.org/10.1242/jeb.020768.

[18] Farina, W. M., Varjú, D. & Zhou, Y. The regulation of distance to dummy flowers during hovering flight in the hawk moth macroglos-sum stellatarum. Journal of Comparative Physiology A 174, 239–247 (2004).

[19] Davidson, J. D., Vishwakarma, M. & Smith, M. L. Hierarchical approach for comparing collective behavior across scales: Cellular systems to honey bee colonies. Frontiers in Ecology and Evolution 9 (2021). URL https://www.frontiersin.org/article/10.3389/fevo.2021.581222. https://doi.org/10.3389/fevo.2021.581222.

[20] Lemanski, N. J., Cook, C. N., Smith, B. H. & Pinter-Wollman, N. A multiscale review of behavioral variation in collective foraging behavior in honey bees. Insects 10 (11), 370 (2019). URL https://pubmed.ncbi.nlm.nih.gov/31731405. https://doi.org/10.3390/insects10110370, 31731405[pmid].

[21] Parrish, J. K. & Edelstein-Keshet, L. Complexity, pattern, and evolutionary trade-offs in animal aggregation. Science 284 (5411), 99–101 (1999). https://doi.org/10.1126/science.284.5411.99.

[22] Seeley, T. The Five Habits of Highly Effective Honeybees (and What We Can Learn from Them): From “Honeybee Democracy” (Princeton University Press, 2010). URL https://books.google.com/books?id=N7KVtAEACAAJ.

[23] Islam, M. S. & Faruque, I. A. Experimental identification of individual insect visual tracking delays in free flight and their effects on visual swarm patterns. bioRxiv (2022). https://doi.org/10.1101/2022.04.06.487367.

[24] Himakalasa, A. & Wongkaew, S. Stability analysis of swarming model with time delays. Advances in Difference Equations 2021 (1), 217 (2021). https://doi.org/10.1186/s13662-021-03379-9.

[25] Svoboda, T., Martinec, D. & Pajdla, T. A convenient multicamera self-calibration for virtual environments. Presence 14 (4), 407–422 (2005). https://doi.org/10.1162/105474605774785325.

[26] Ljung, L. System Identification: Theory for the User Prentice Hall information and system sciences series (Prentice Hall PTR, 1999). URL https://books.google.com/books?id=nHFoQgAACAAJ.

[27] Bennet, D. J. & McInnes, C. R. Distributed control of multi-robot systems using bifurcating potential fields. Robotics and Autonomous Systems 58 (3), 256–264 (2010). https://doi.org/https://doi.org/10.1016/j.robot.2009.08.004, towards Autonomous Robotic Systems 2009: Intelligent, Autonomous Robotics in the UK.

[28] Lindley, B., Mier-Y-Teran-Romero, L. & Schwartz, I. B. Randomly distributed delayed communication and coherent swarm patterns. IEEE Int Conf Robot Autom (2012).

[29] Bernard, S., Bélair, J. & Mackey, M. Sufficient conditions for stability of linear differential equations with distributed delay. Discrete and Continuous Dynamical Systems. Series B (2001). https://doi.org/10.3934/dcdsb.2001.1.233.

